# CRISPR-Cas9/Cas12a Systems for efficient genome editing and large genomic fragment deletions in *Aspergillus niger*

**DOI:** 10.1101/2024.06.24.600459

**Authors:** Guoliang Yuan, Shuang Deng, Jeffrey J. Czajka, Ziyu Dai, Beth A. Hofstad, Joonhoon Kim, Kyle R. Pomraning

**Author notes:** Corresponding authors: Guoliang Yuan; Kyle R. Pomraning.

## Abstract

CRISPR technology has revolutionized fungal genetic engineering by accelerating the pace and expanding the feasible scope of experiments in this field. Among various CRISPR-Cas systems, Cas9 and Cas12a are widely used in genetic and metabolic engineering. In filamentous fungi, both Cas9 and Cas12a have been utilized as CRISPR nucleases. In this work we first compared efficacies and types of genetic edits for CRISPR-Cas9 and -Cas12a systems at the polyketide synthase (*albA*) gene locus in *Aspergillus niger*. By employing a tRNA-based gRNA polycistronic cassette, both Cas9 and Cas12a have demonstrated remarkable editing efficacy. Cas12a demonstrated superiority over Cas9 protein when one gRNA was used for targeting, achieving an editing efficiency of 89.5% compared to 15% for Cas9. Moreover, when employing two gRNAs for targeting, both systems achieved up to 100% editing efficiency for single gene editing. In addition, the CRISPR-Cas9 system has been reported to induce large genomic deletions in various species. However, its use for engineering large chromosomal segments deletions in filamentous fungi still requires optimization. Here, we engineered Cas9 and - Cas12a-induced large genomic fragment deletions by targeting various genomic regions of *A*. *niger* ranging from 3.5 kb to 40 kb. Our findings demonstrate that targeted engineering of large chromosomal segments can be achieved, with deletions of up to 66.7% efficiency. Furthermore, by targeting a secondary metabolite gene cluster, we show that fragments over 100 kb can be efficiently and specifically deleted using the CRISPR-Cas9 or -Cas12a system. Overall, in this paper, we present an efficient multi-gRNA genome editing system utilizing Cas9 or Cas12a that enables highly efficient targeted editing of genes and large chromosomal regions in *A*. *niger*.

## 1 Introduction

Filamentous fungi play a vital role in biotechnology as they serve as indispensable hosts for producing a wide arrange of compounds essential for various industries. They are widely utilized in the production of enzymes crucial for food processing, pharmaceuticals, commodity chemicals, organic acids, and biofuels (Füting et al. 2021; Lübeck and Lübeck 2022). Their remarkable metabolic diversity and ability to thrive in diverse environments have made fungi indispensable workhorses for large-scale fermentations, and fungi have helped drive innovation in biotechnological applications within industrial processes. Employing genetic engineering to understand and modify filamentous fungi is pivotal for enhancing productivity and creating customized strains for diverse biotechnological applications. However, genetic modifications in fungi have been significantly hampered by the scarcity of genetic tools, with traditional genome editing predominantly relying on low-efficiency homologous recombination (HR) and constrained selectable marker availability (Nødvig et al. 2015). Hence, there is a growing demand for the development of versatile methods that enable more efficient, easier, and flexible genetic manipulation of filamentous fungi.

The development of the clustered regulatory interspaced short palindromic repeats/CRISPR-associated nucleases (CRISPR-Cas) genome editing systems have significantly improved the efficiency of introducing desired mutations into the genome compared to previous genome-editing tools, thereby accelerating both fundamental research and the application of fungi in industrial biotechnology (Cong et al. 2013; Ouedraogo and Tsang 2020). The CRISPR-Cas system forms a complex with a guide RNA (gRNA), directing the nuclease to the target DNA. The type II CRISPR-Cas9 system, derived from *Streptococcus pyogenes* (SpCas9), is among the most extensively studied and widely utilized categories of CRISPR-Cas systems due to their efficiency and simplicity in genome editing (Adli 2018). The Cas9 induces DNA double-strand breaks at the target site, which can undergo repair via either the non-homologous end-joining pathway (NHEJ) or homology-directed repair (HDR), resulting in genome editing. Since 2015, CRISPR/Cas9-based genome editing of filamentous fungi has been carried out in several fungal species. For example, CRISPR-Cas9 system has been employed to introduce mutations into single loci in six different *Aspergillus* species, such as *A*. *nigulans*, *A*. *niger*, *A*. *aculeatus*, *A*. *luchuensis*, *A*. *brasiliensis*, and *A*. *carbonarius* (Nødvig et al. 2015). CRISPR-Cas9-based editing methods have also been effectively applied in fungal species such as *A*. *orzyae*, *Trichoderma reesei*, and *Nodulisporium* sp., demonstrating the versatility and wide-ranging applicability of this approach (Liu et al. 2015; Katayama et al. 2016; Zheng et al. 2017). However, CRISPR- Cas9 relies on recognizing a specific protospacer adjacent motif (PAM) sequence NGG, which limits the genomic sites available for targeting editing. This constraint poses a specific challenge when attempting to edit A/T base pair rich regions of the genome. Consequently, this requirement restricts our capacity to target certain sequences with the CRISPR system, prompting widespread endeavors either to discover alternative PAM sequences or to relax the PAM requirement. CRISPR-Cas12a (also known as Cpf1), a Class II type V endonuclease, is a novel RNA-guided enzyme that has recently emerged as an alternative tool for genome editing (Fonfara et al. 2016). Unlike Cas9, which targets guanine (G)-rich sequences, Cas12a identifies specific thymine (T)-rich PAMs, which greatly expands the number of possible editing sites beyond those targeted by Cas9 (Kim et al. 2020). Recently, various research groups have successfully established functional CRISPR-Cas12a systems in various filamentous fungi, such as *A*. *niger*, *A*. *oryzae*, *A*. *sojae*, *A*. *aculeatus*, and *A*. *nidulans* (Vanegas et al. 2019; Abdulrachman et al. 2021; Katayama and Maruyama 2022).

In addition to single gene deletions or edits, generating large chromosomal deletions are crucial for both functional genomics, genome reduction, and organism breeding, particularly in microorganisms used in industrial processes (Takahashi et al. 2008). These deletions help mitigate potential risks associated with pathogenic or toxic genes that may be expressed under specific conditions. Although methods for inducing chromosomal deletions in both bacteria and *Saccharomyces cerevisiae* have been intensively studied, further exploration is needed to apply these techniques to delete large chromosomal segments in filamentous fungi (Hegemann et al. 2014; Standage-Beier et al. 2015). However, establishing a general technique for removing unfavorable genomic regions from industrially important microorganisms is desirable but challenged by low homologous recombination frequencies, especially in wild-type strains. The CRISPR/Cas9 system, employing dual sgRNA-directed large gene deletion, is a robust tool for large-scale chromosomal deletions. Recent studies have demonstrated CRISPR/Cas9-induced large genomic deletions ranging from 20 kb to 5 Mb in *Caenorhabditis elegans*, maize, rabbit, human cell lines, and mouse (Chen et al. 2014; Essletzbichler et al. 2014; Zhou et al. 2014; Song et al. 2016; Kato et al. 2017; Li et al. 2023). However, it is noted that efficiency decreases as the size of the deleted fragment increases.

Developing highly efficient genome editing systems is crucial for metabolic engineering, particularly for the refining and customization of industrial strains. These processes often entail multiple rounds of modification, highlighting the necessity for precise and effective genome editing tools. In this study, we introduce two vector-based CRISPR platforms, CRISPR-Cas9 and CRISPR-Cas12a, designed for highly efficient fungal CRISPR-mediated gene editing. Our findings demonstrate the versatility of both Cas9 and Cas12a systems in *A*. *niger*. Moreover, we showcase their efficacy in inducing large chromosomal segment deletions with up to 102 kb, further broadening their utility in genetic manipulation. These advancements offer promising avenues for accelerating strain optimization and advancing metabolic engineering strategies in industrial biotechnology.

## 2 Materials and methods

### 2.1 Strains and culture conditions

*Escherichia coli* strain DH5α was used as the host for routine cloning procedures. *A*. *niger* wild-type strain ATCC 11414 from the American Type Culture Collection (Rockville, MD, 20852) was cultivated on complete medium (CM) plates at 30°C for culture maintenance and spore preparation. Cultures on CM agar plates were incubated at 30°C for 4 days, followed by harvesting of spores through washing with sterile 0.5% Tween 80. Spore counting was performed using a hemocytometer. Both CM and minimal medium (MM) were prepared according to the method described by Bennett and Lasure (Bennett and Lasure 1991).

### 2.2 Vector construction

A total of 18 vectors were utilized in this study (Table 1). The pFC332 vector (Plasmid #87845), obtained from Addgene, served as an empty vector for cloning purposes. Initially, pFC332 was linearized using the PacI restriction enzyme. Subsequently, the pGY5 vector was constructed by assembling the linearized pFC332 with a gBlocks Gene Fragment 7_gblocks via HiFi DNA assembly. Similarly, the pGY6 vector was generated by assembling the linearized pFC332 with two gBlocks Gene Fragments, 16_gblocks and 17_gblocks, using HiFi DNA assembly. pGY18 was assembled by combining the linearized pFC332 (via BsaI) with three PCR products, namely PCR8768, PCR6926, and PCR7086, using HiFi DNA assembly. pGY53 was constructed by combining the linearized pGY18 (via BsaI and BamHI) with two PCR products (PCR162216 and PCR165186), along with three gBlocks Gene Fragments (77_gblocks, 78_gblocks, and 79_gblocks), using HiFi DNA assembly. pGY54 was created by combining the linearized pGY53 (via BsaI) with a gBlocks Gene Fragment 80_gblocks using NEBridge Golden Gate Assembly Kit (BsaI-HF v2). Similarly, pGY55 was assembled by combining the linearized pGY53 (via BsaI) with two gBlocks Gene Fragments, 81_gblocks and 82_gblocks, also utilizing Golden Gate assembly. pGY42, pGY56, pGY76, pGY77, pGY108, pGY109, and pGY177 were all generated by ligating linearized pGY18 (via BsaI) with their respective gBlocks Gene Fragments using Golden Gate assembly. Similarly, pGY158, pGY175, pGY176, and pGY178 were created by ligating linearized pGY53 (via BsaI) with their corresponding gBlocks Gene Fragments using Golden Gate assembly. Q5 Master Mix was employed for all cloning PCR according to the manufacturer’s instructions. All cloning enzymes and restriction enzymes utilized in this study were sourced from New England Biolabs (Ipswich, MA, USA). All plasmids were verified by Sanger sequencing. Additionally, all gBlocks Gene Fragments were acquired from IDT (Supplementary Table S1). For experimental vectors, the empty vector for Cas9, designated as pGY18, and the empty vector for Cas12a, known as pGY53, will be made accessible through Addgene.

**Table 1.**
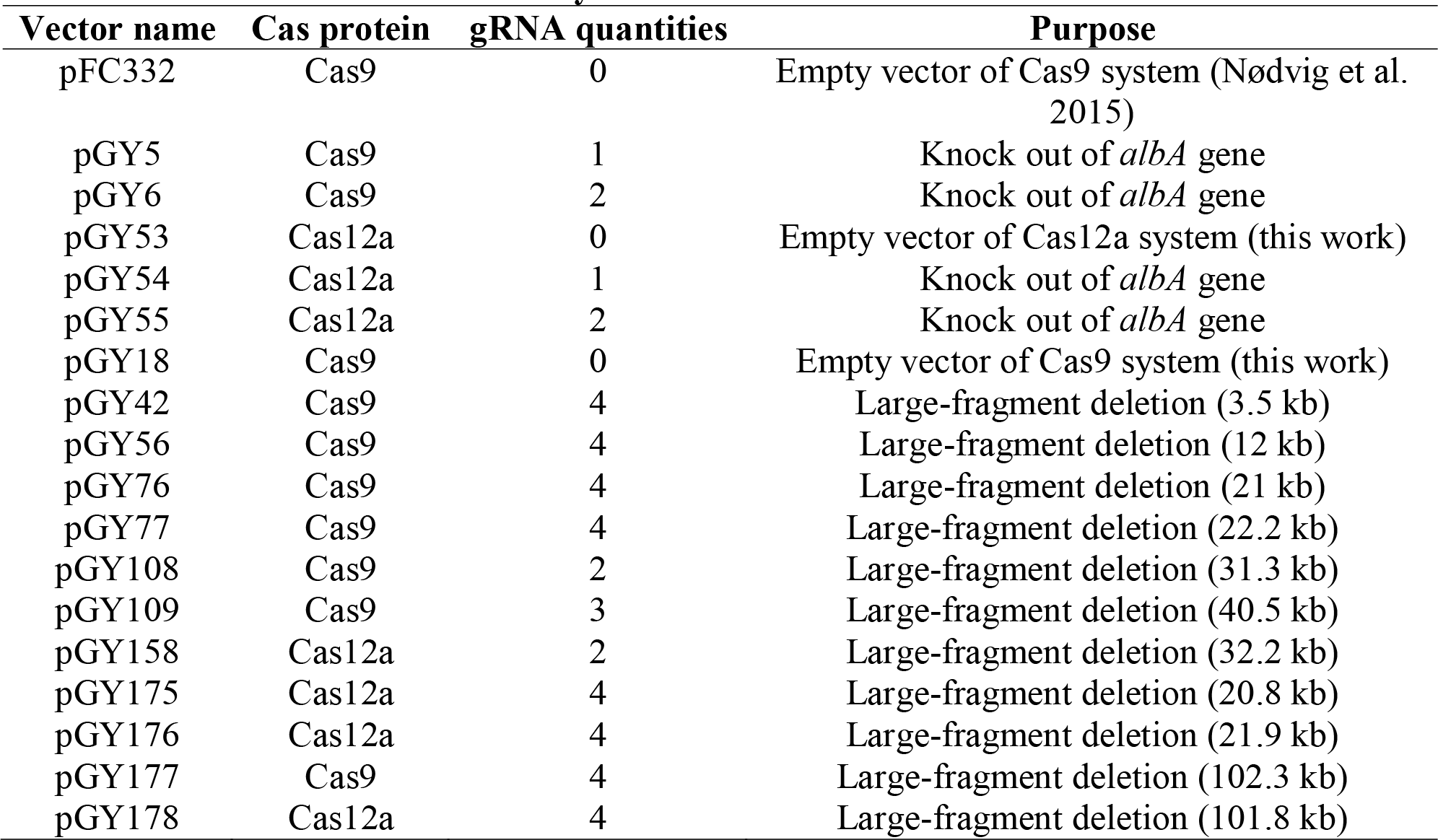
All vectors used in this study.

### 2.3 Design of gRNAs

All gRNAs were designed using the online tool CHOPCHOP (Labun et al. 2019). A total of 35 gRNAs were utilized in this study (Table 2).

**Table 2.**
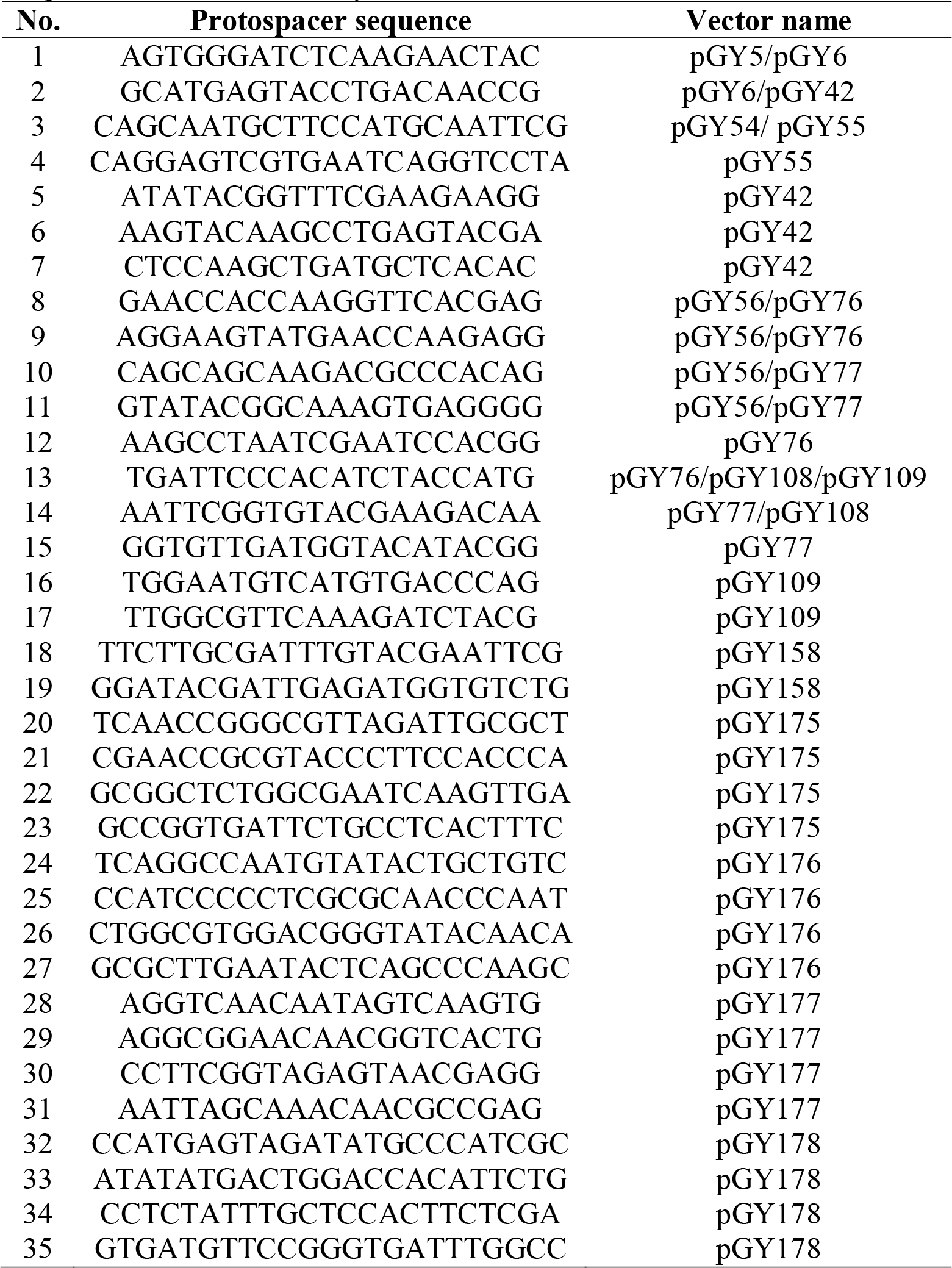
All gRNAs used in this study.

### 2.4 Protoplast transformation

A standard polyethylene glycol (PEG)-mediated protoplast transformation protocol was employed in *A*. *niger as previously described*. About 1-2 µg of plasmid in a volume of 10 µL or less was added to 100 µL of protoplasts in 15 ml centrifuge tube, which was gently mixed by tapping the tube and incubated on ice for 15 minutes. Next, 1mL of 40% PEG was added, gently mixed by taping the tube, and kept at room temperature for 15 minutes. Following this, 5 mL of minimum medium containing 1M Sorbitol was added and the tube was placed horizontally in the incubator shaker and shaken gently (∼80 RPM) at 30°C for 1 hour. Subsequently, the protoplasts were pelleted down by centrifugation at 800 g and 4°C for 5 minutes. The pelleted protoplasts were resuspended in 15 mL of minimal medium with 1M sorbitol and 0.8% agar. A 300µg/mL of hygromycin or geneticin (G418) were used for antibiotic selection in *A. niger*. Finally, the resuspended cells with proper antibiotics were poured onto a petri dish and culture at 30°C for 1∼2 days.

### 2.5 Single colony isolation

To quantify the editing percentage of CRISPR, single colonies were isolated under a dissection microscope (Leica MZ16) from the transformation plate after 19-24 hours post-transformation. Each colony was transferred to a slant containing 1.5 mL of MM medium supplemented with 0.8% agar and 200µg/mL hygromycin or G418. The slants were then cultured at 30°C until spores were observed.

### 2.6 White spore isolation

To isolate the white spores from slants containing both white and black spores, a syringe was employed under a dissection microscope to carefully collect the white spores. Subsequently, these collected white spores were inoculated onto fresh CM slants to undergo purification for further analysis and experimentation.

### 2.7 PCR genotyping

The spores were collected by rinsing with sterile 0.5% Tween 80. Subsequently, these harvested spores were directly utilized for Squash-PCR genotyping following previously established procedures (Yuan et al. 2023). Genotyping PCR was performed using GoTaq Green Master Mixes (Promega). For each reaction, 1 μL of squashed spore solution was used as a template in a total PCR reaction volume of 20 μL. The PCR cycling conditions were as follows: initial denaturation at 95 °C for 2 min, followed by 35 cycles of denaturation at 95 °C for 30 s, annealing at 55 °C for 30 s, and extension at 72 °C for 1 min, with a final elongation step at 72°C for 5 min. All DNA oligos used for genotyping PCR are listed in Supplementary Table S2.

### 2.8 Sanger sequencing

PCR products were purified using the QIAquick gel extraction kit from QIAGEN N.V. (Venlo, Netherlands) and subsequently sequenced by GENEWIZ (South Plainfield, NJ, USA). The DNA oligos used for genotyping PCR are additionally employed for Sanger sequencing.

## 3 Results

### 3.1 Single gene editing in *A*. *niger* mediated by CRISPR-Cas9 and Cas12a

CRISPR genome editing tools are capable of editing the genome at both single and multiple locations. To enhance the efficiency of CRISPR gene editing systems, we utilized a tRNA-based gRNA expression strategy to boost the expression of multiple gRNAs. In this method, individual gRNAs are released by the cellular endogenous tRNA processing machinery, a mechanism that has been demonstrated to work efficiently across multiple species (Čermák et al. 2017; Song et al. 2019; Zhang et al. 2019). Since both the Cas9 and Cas12a systems have been established in *A*. *niger*, our objective is to compare the editing efficiency of these two systems by targeting the *albA* gene . This gene is a classical reporter gene with white *A. niger* spores as marker. (Vanegas et al. 2019; Li et al. 2021). Additionally, we aim to characterize single gene editing mediated by both single gRNA and double gRNAs to compare editing efficiency and provide practical guidance for their use. The *tef1* promoter and its terminator were employed to control the expression of Cas9 or Cas12a in all vectors, while a U3 promoter and its terminator derived from *Aspergillus fumigatus* were utilized to drive the expression of the tRNA-gRNA array (Figure 1A). Recently, it has been demonstrated that among the endogenous tRNAs in *A*. *niger*, tRNA^Ala^, tRNA^Phe^, tRNA^Arg^, tRNA^Ile^, and tRNA^Leu^ are particularly effective in enhancing gRNA release compared to others (Li et al. 2021). Therefore, we applied these tRNA sequences to boost the expression and release of gRNAs in both Cas9 and Cas12a systems. For the Cas9 systems, single and double gRNAs targeting the coding region of *albA* were integrated into vectors pGY5 and pGY6, respectively, as described in Method 2.2 (Figure 1A). However, the original empty vector of Cas9 (pFC332) lacks the U3 promoter and terminator necessary for the expression of gRNAs. Additionally, its lack of suitable restriction enzyme sites makes ligation with multiple gRNA fragments challenging. Therefore, we modified it by incorporating the U3 promoter and terminator. Furthermore, we adapted this vector to pGY18 for BsaI-based Golden Gate assembly, allowing efficient ligation of multiple gRNA fragments, as demonstrated by its capability to tandemly assemble 1 to 16 gRNAs in a one-pot reaction (Yuan et al. 2022). By replacing Cas9 with Cas12a in pGY18, we further engineered a novel Cas12a vector, pGY53, designed for the ligation of multiple gRNA fragments through Golden Gate assembly. Using pGY53 as the backbone, we generated vectors pGY54 and pGY55, which contain single and double gRNAs, respectively, following the procedures described in Method 2.2 (Figure 1A).

**Figure 1.**
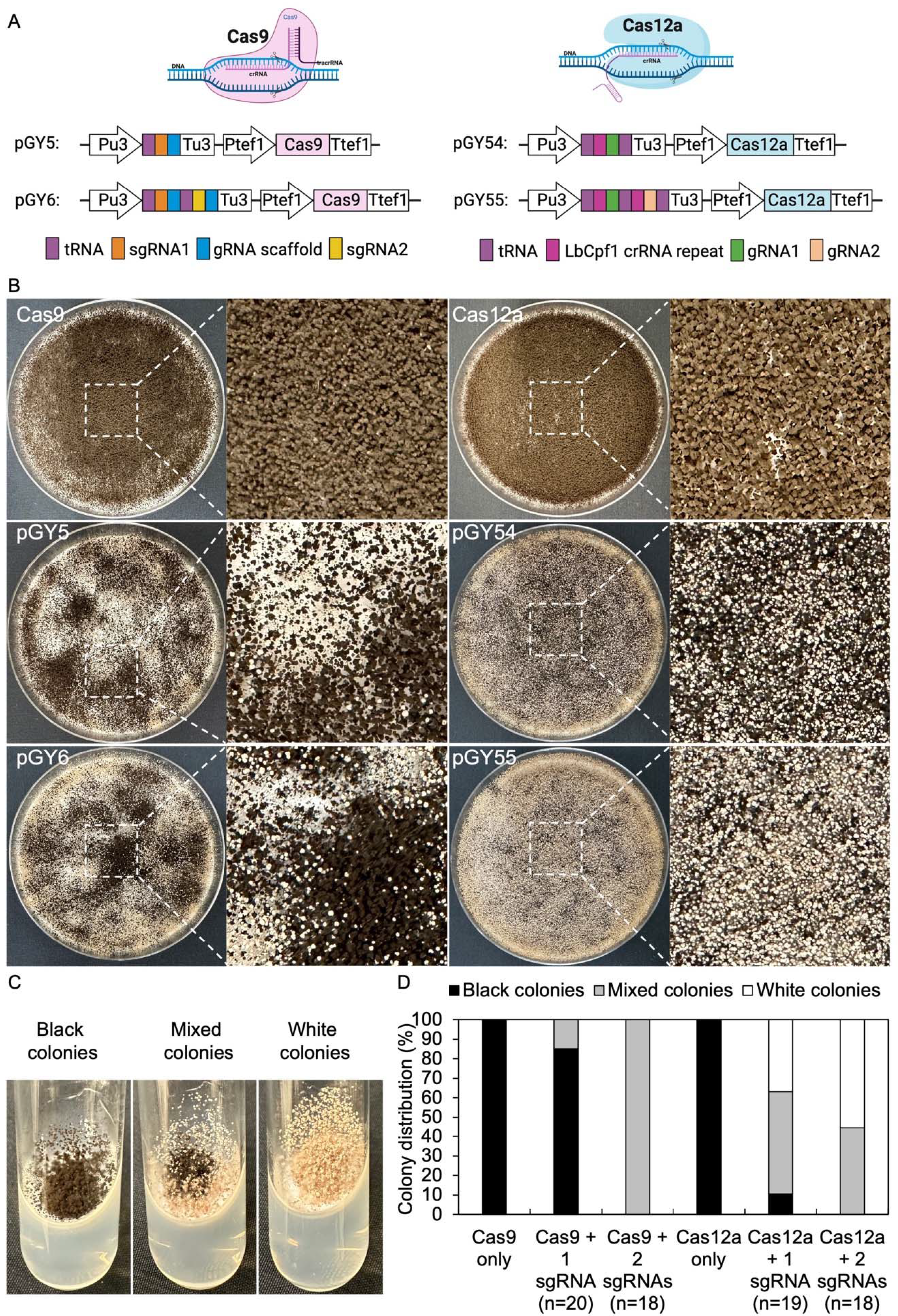
CRISPR-Cas9 and Cas12a mediated single gene editing in *A*. *niger*. (A). Illustration of CRISPR-Cas9 and Cas12a constructs. (B). Phenotypic effects of *albA* mutations induced by Cas9 and Cas12a systems. C. Phenotypic characteristics of *albA* mutants following single colony isolation on an agar slant. D. Editing efficacy comparison between CRISPR-Cas9 and Cas12a systems.

Next, we conducted protoplast transformation as described in Method 2.4 to transform vectors pGY5, pGY6, pGY54, and pGY55 into *A*. *niger* strain 11414, with pFC332 and pGY53 serving as the empty vector controls. Due to the high transformation efficiency, tens to hundreds of colonies were observed after transformation, resulting in the entire plate being covered with hyphae and spores (Figure 1B). Pure black spores were observed on the plates of pFC332 (Cas9- only) and pGY 53 (Cas12a-only), while large quantities of white spores were present on all plates of pGY5, pGY6, pGY54, and pGY55, indicating successful genome editing events (Figure 1B). More specifically, the abundance of white spores on plates of pGY6, pGY54, and pGY55 is noticeably higher than that of pGY5, suggesting a significant increase in editing efficiency. To quantify the editing percentage, we performed single colony isolation following the procedure outlined in Method 2.5. After single colony isolation, three distinct phenotypes were observed: those containing pure black spores, pure white spores, and mixed black and white spores (Figure 1C and Supplementary Figure S1). It was evident that slants containing pure white spores and mixed spores represented genome-edited events. After quantification, we observed editing efficiencies of 15% and 89.5% for Cas9 and Cas12a, respectively, when single gRNA was used to target the *albA*. Remarkably, both editing efficiencies increased to 100% when double gRNAs were used to target the *albA* (Figure 1D). It should be noted that no events of pure white spores were observed in the Cas9 systems. In contrast, white spores appeared in 36.8% and 55.6% of the cases in the Cas12a system when single and double gRNAs were used, respectively (Figure 1D and Supplementary Figure 1). This difference underscores the distinct phenotypic outcomes associated with Cas9 and Cas12a editing systems. To ensure the purity of editing events, white spores were isolated according to the procedure outlined in Method 2.6. Following this, we performed Squash-PCR genotyping and subsequent Sanger sequencing to validate the purified editing events, providing further confirmation of the desired genomic modifications. In the genotyping PCR, three selected events of pGY5 exhibited bands similar to the positive control, while 4 out of 5 tested events of pGY6 displayed lower bands compared to the wild-type band, indicating the occurrence of deletions in the targeted region (Figure 2A). Notably, in event #14 of pGY6, double bands were observed, indicating the mixture of both wild-type and mutant. The PCR genotyping was thoroughly validated by Sanger sequencing. For example, small deletions of 1 bp and 18 bp were observed in the editing events of pGY5, explaining that there were no obvious size differences between the wild type and mutants due to these small deletions (Figure 2A and B). In contrast, larger deletions ranging from 90 to 125 bp were precisely detected between the two applied gRNAs, corroborating the lower mutant bands observed in the PCR genotyping (Figure 2A and B). In the testing of Cas12a system, five editing events from pGY54 and pGY55, respectively were selected for PCR genotyping. Surprisingly, six out of the total ten tested events failed to yield any PCR products. This could be attributed to either low-quality DNA templates or the partial or complete removal of the amplification region by CRISPR. Notably, the same DNA templates successfully amplified PCR products using other primer pairs, suggesting that the latter is the primary reason for the lack of amplification. Large deletions, such as 750 and 127 bp, were observed in event #6 of pGY54 and event #5 of pGY55, respectively (Figure 2B). This observation is completely consistent with the mutant bands detected in the genotyping PCR (Figure 2A). Interestingly, a 17 bp deletion and a 4 bp deletion were detected in event #4 of pGY55, corresponding to the sites targeted by the first and second gRNAs, respectively (Figure 2B). However, the sequence between the two gRNAs was not removed, indicating that the editing of these two sites likely did not occur simultaneously.

**Figure 2.**
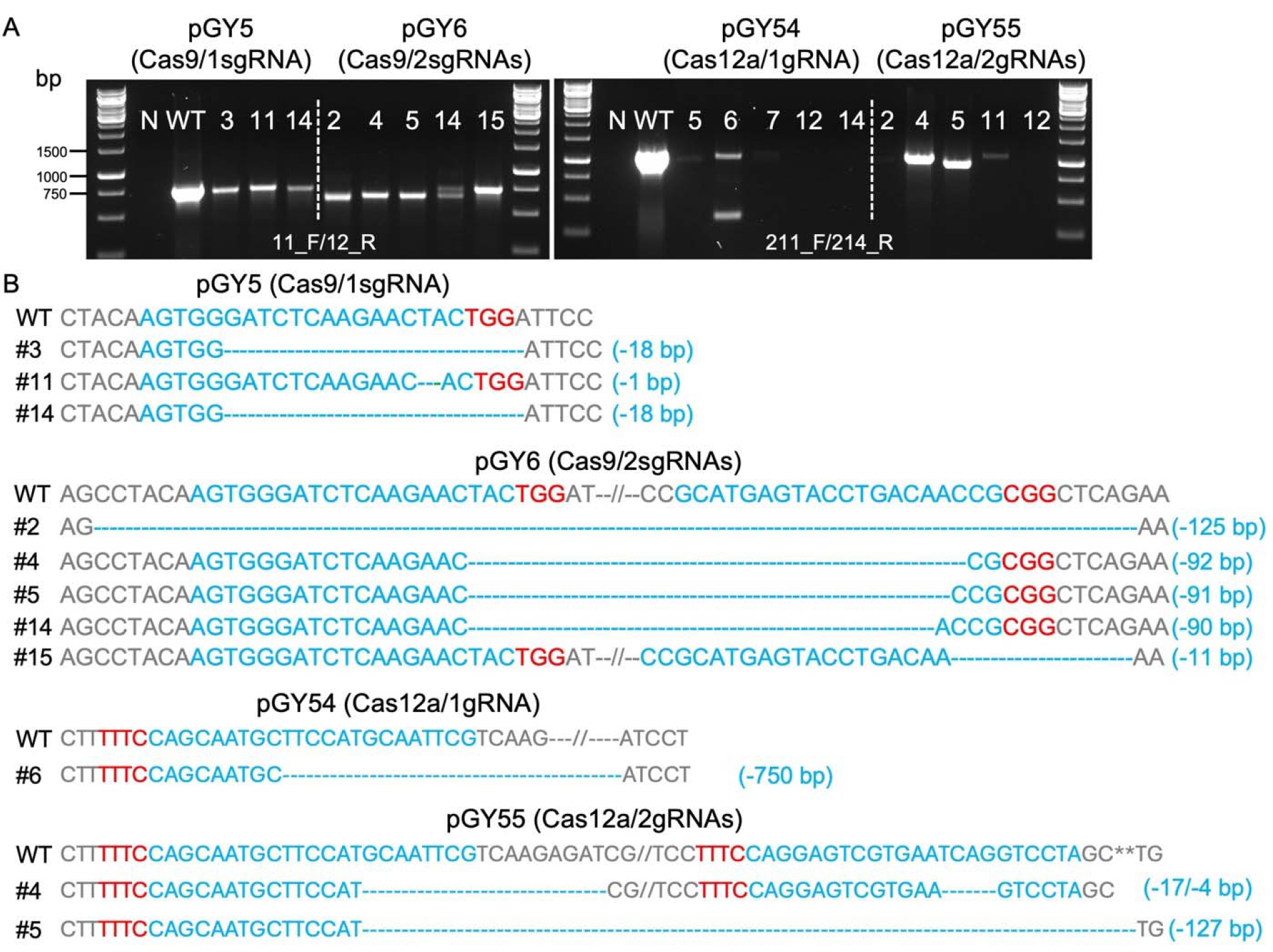
Analysis of editing events mediated by Cas9 and Cas12a systems. (A). PCR genotyping for mutant screening. N, water as negative control; WT, wild type as positive control. (B). Sanger sequencing for mutation confirmation. PAM sequences are highlighted in red, while gRNA sequences are highlighted in blue.

Overall, both Cas9 and Cas12a demonstrated exceptionally high efficiency for single gene editing, particularly when double gRNAs were applied in *A*. *niger*. Cas12a exhibited superiority over the Cas9 system, as it showed significantly higher editing efficiency than Cas9 when only one gRNA was used for gene editing. Furthermore, achieving pure mutants seemed to be significantly easier with Cas12a after transformation, thereby facilitating the single spore isolation required for the strain production test.

### 3.2 Large chromosome segment deletion mediated by CRISPR-Cas9 and Cas12a

To evaluate the capacity for targeted CRISPR-mediated large-scale genomic deletions in *A*. *niger*, we selected both the upstream and downstream sequences of the *albA* gene as the target region (Figure 3A). Multiple regions were selected according to the anticipated size of deletions, ranging incrementally from smaller to larger: 3.5 kb, 10 kb, 20 kb, 30 kb, and 40 kb, and diversified the selection to include a broad array of deletion sizes within this range (Figure 3A). To eliminate a long DNA sequence, a double-strand break (DSB) needed to be induced both upstream and downstream of the target region. Therefore, we designed gRNA1 and gRNA2 to target the upstream and downstream regions of the target locus, respectively, with high targeting efficiency as predicted by CHOPCHOP (Labun et al. 2019) (Figure 3B). In case these gRNAs did not exhibit sufficient efficiency, we additionally designed gRNA3 and gRNA4 proximal to gRNA1 and gRNA2, respectively, to ensure comprehensive coverage. Technically, target regions exceeding 10 kb pose challenges for straightforward amplification via standard PCR methods. Upon complete removal of the target region, amplifying the edited segment would be facilitated due to shorter amplification lengths with external primer pairs, while the internal sequence of the target regions should remain undetectable by PCR when using internal primer pairs (Figure 3B).

**Figure 3.**
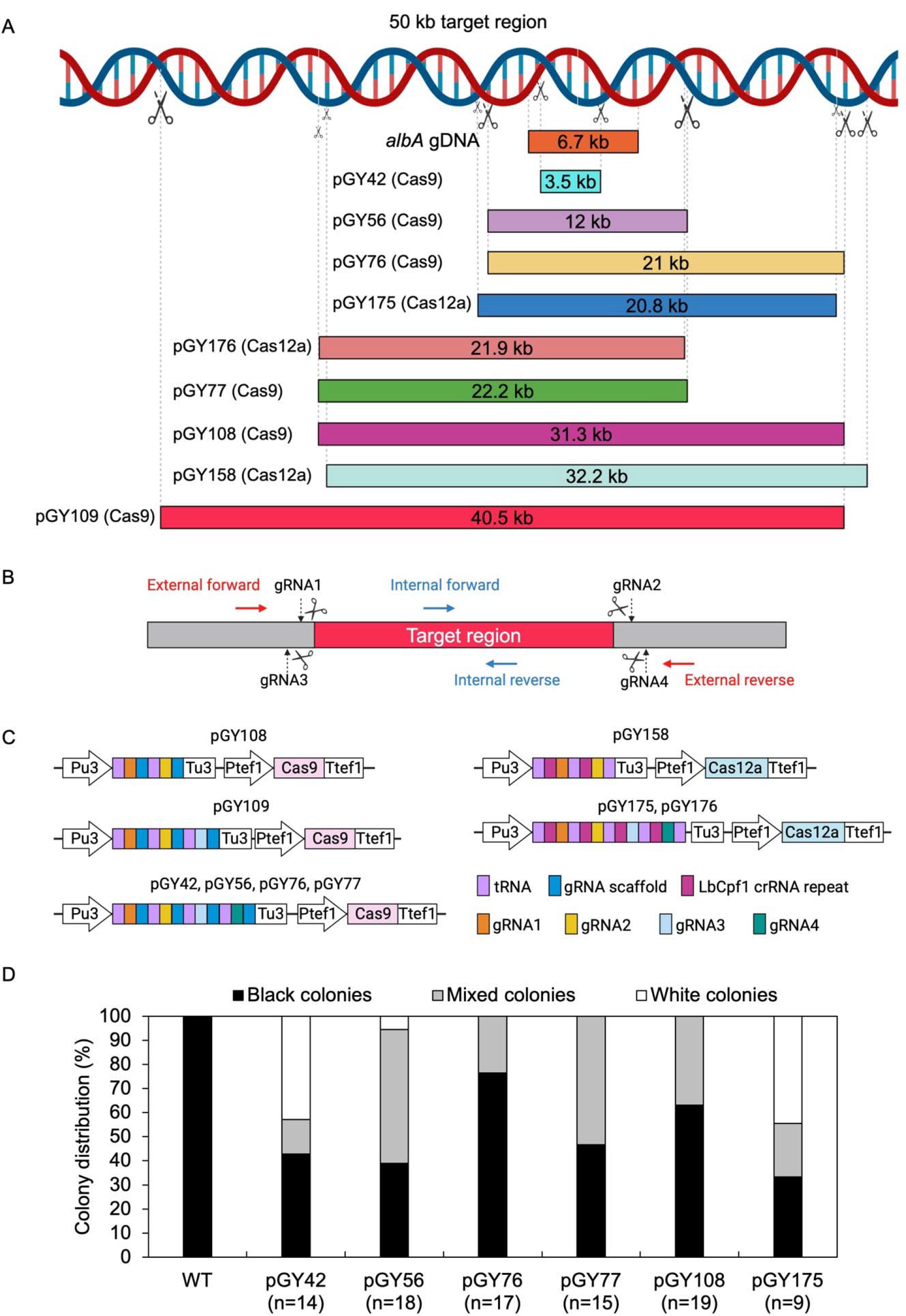
CRISPR-Cas9 and Cas12a mediated large chromosome segment deletion in *A*. *niger*. (A). Selected target region for large fragment deletion. (B). gRNA and primer design for large fragment deletion. (C). Illustration of CRISPR-Cas9 and Cas12a constructs. (D). Editing efficiency of large fragment deletion. Note: Insufficient data for the analysis of colony distribution with pGY109, pGY158, or pGY176 construct due to the failure of isolated colony growth.

Based on above principles, multiple gRNAs were designed for both Cas9 and Cas12a systems. For instance, four gRNAs were utilized for vectors pGY42, pGY56, pGY76, pGY77, pGY175, and pGY176; three gRNAs were employed for vector pGY109, while two gRNAs were utilized for vectors pGY108 and pGY158 (Figure 3A and C).

Following protoplast transformation, all plates containing nine different CRISPR constructs exhibited white spores, contrasting with the black spores observed on the wild-type (WT) plate, indicating successful elimination of the target region containing the *albA* gene (Figure 4). The quantity of white spores significantly varies across the plates, with notably higher abundance observed in those of pGY42, pGY175, and pGY176 compared to others, suggesting a higher editing efficiency in these plates (Figure 4). Due to the dense coverage of hyphae and spores on the plates hindering direct editing percentage calculation, we conducted single colony isolation for pGY42, pGY56, pGY76, pGY77, pGY108, and pGY175 at the early post-transformation stage, following Method 2.5. We observed similar phenotypes across plates, including those with pure black spores, pure white spores, and mixed black and white spores, as illustrated in Figure 1C and Supplementary Figure S2. Following quantification, we determined that the editing percentage of all constructs ranged from 23.5% to 66.7%. Notably, four constructs demonstrated editing efficiency exceeding 50%, with percentages of 57.1%, 61.1%, 53.3%, and 66.7% for pGY42, pGY56, pGY77, and pGY175, respectively (Figure 3D).

**Figure 4.**
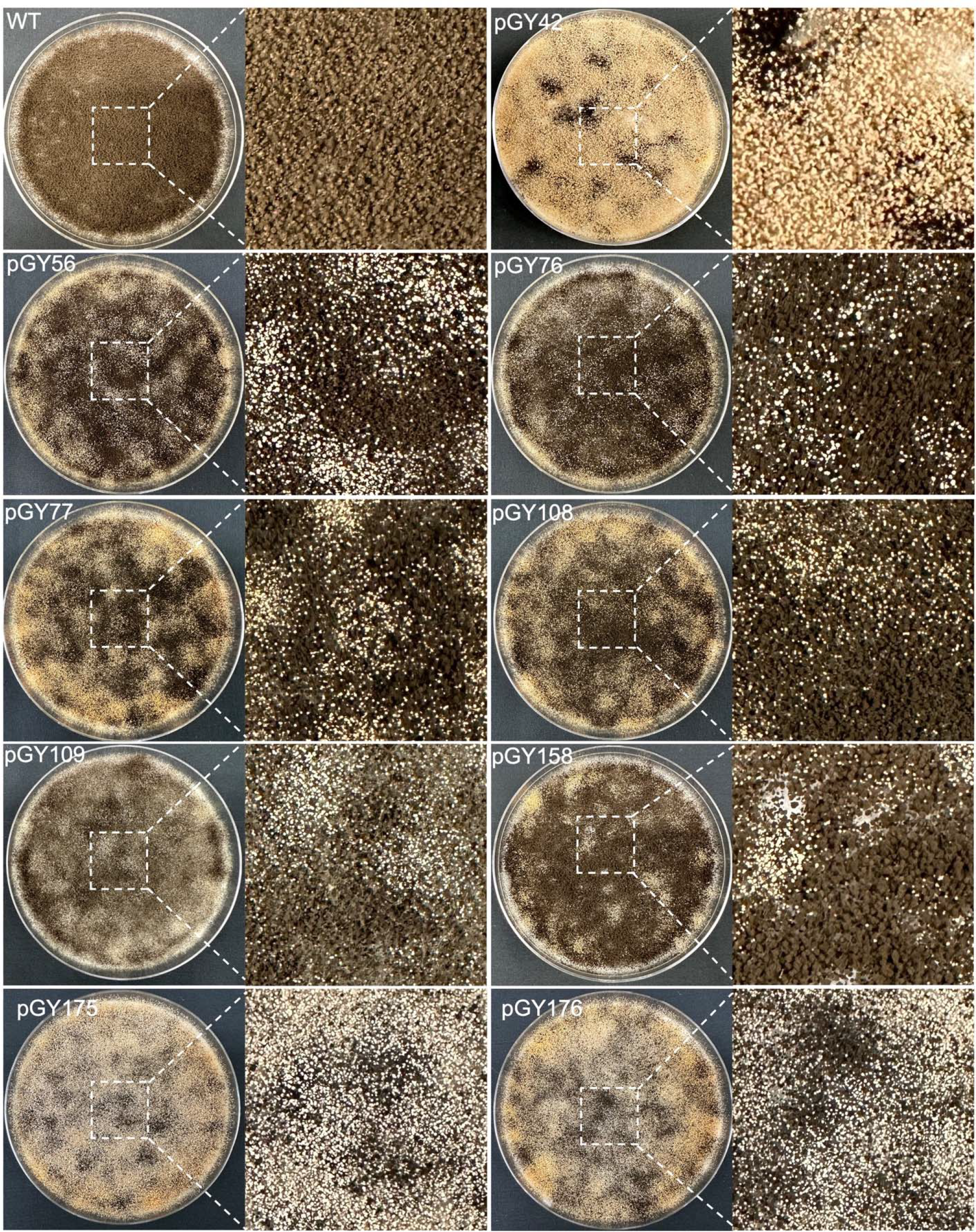
Phenotypic effects of large chromosome fragment deletion induced by CRISPR- Cas9 and Cas12a systems.

After isolation of the white spores was performed (Method 2.6), PCR genotyping and Sanger sequencing were employed to validate and determine the precise sequence and size of the large fragment deletions. As depicted in Figure 3B, external primers targeting various regions were utilized to amplify the full length of the target region. Concurrently, identical internal primers were used to amplify the internal sequence of the target region (located within the *albA* gene) for all constructs, except for pGY42, where the target region is situated within the *albA* gene itself. For instance, external primers 211_F/212_R were specifically designed to amplify the entire target region of pGY42, which spans 4,924 bp in length. Conversely, three edited events, #5, #14, and #15, yielded PCR products ranging in size from approximately 1.3 to 1.6 kb. This indicates that fragments ranging from approximately 3.3 to 3.6 kb in length were deleted from the target region. The external primers 224_F/225_R were used to amplify the complete target region of pGY56, spanning a length of 12,455 bp. However, events #3 and #5 produced 1 kb PCR products only, and the internal region (*albA* gene) was not detected using internal primers 211_F/214_R, indicating the elimination of a sequence 11 kb in length (Figure 5A). Indeed, the sequence spanning 11,506 bp between the two gRNAs was deleted in event #3, as confirmed by Sanger sequencing (Figure 5B). Using similar strategies, we investigated all the rest of the constructs. Due to variations in the target region, the full length of target regions ranged from approximately 21 kb to 42 kb. However, all the tested editing events yielded PCR products of less than 1.5 kb, and no obvious internal regions were detected (Figure 5A). This finding indicates deletions occurring within a range from as small as 20 kb to as large as 40 kb in these events. Interestingly, the deleted sequences and their sizes were accurately determined by Sanger sequencing, perfectly matching with the corresponding PCR genotyping results. For example, a 21,100 bp deletion was identified in event #1 of pGY76, a 22,157 bp deletion in event #2 of pGY77, a 31,311 bp deletion in event #1 of pGY108, a 40,489 bp deletion in event #2 of pGY109, a 32,141 bp deletion in event #2 of pGY158, a 20,750 bp deletion in event #19 of pGY175, and a 21,873 bp deletion in event #11 of pGY176 (Figure 5A and B). In addition, for those constructs employing two gRNAs to target upstream or downstream sequences, we observed deletions occurring at different locations targeted by the respective gRNAs, as demonstrated in constructs pGY76 and pGY176 (Figure 5B). Taken together, our findings demonstrate that large-scale genomic sequences can be efficiently deleted using CRISPR-Cas9 or Cas12a systems, with deletions as large as 40.5 kb achieved in *A*. *niger*.

**Figure 5.**
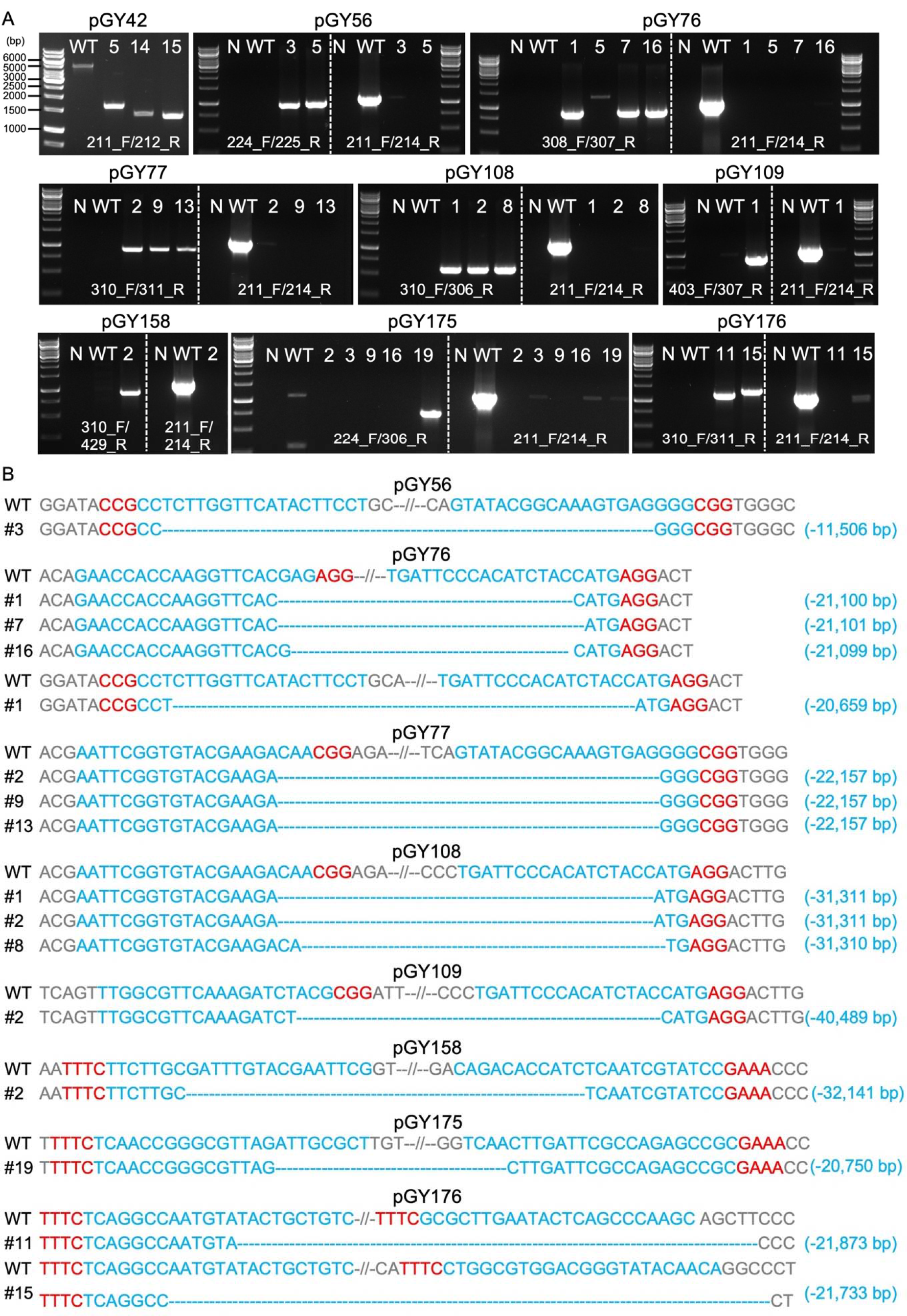
Analysis of large fragment deletion mediated by Cas9 and Cas12a systems. (A). PCR screening for mutants. N, water as negative control; WT, wild-type as positive control. (B). Sanger sequencing for mutation confirmation. PAM sequences are highlighted in red, while gRNA sequences are highlighted in blue.

### 3.3 Deletion of a 100-kb fragment induced by CRISPR-Cas9 and Cas12a

Although we demonstrated deletion of large chromosomal segments up to approximately 40.5 kb within the vicinity of *albA*, we were unable to isolate larger deletions from 50 to 100 kb due to the presence of genes likely to be essential for growth and sporulation. When extending the study to larger regions, the transformation plates yielded no typical white spores with these constructs. To determine the capacity of this technique, we chose an alternative genomic region containing a NRPS (nonribosomal peptide synthase) gene cluster, which involves in synthesis of siderophores such as desmalonichrome and is presumed to be transcriptionally silent under certain cultivation conditions (Mo et al. 2016; Kwon et al. 2021). The NRPS cluster spans a total size of 97,809 bp. To target this region, we designed two vectors: Cas9 vector pGY177 and Cas12a vector pGY178, intended to induce deletions of 102.3 kb and 101.8 kb, respectively (Figure 6A). Following protoplast transformation, 20 colonies were randomly chosen from the transformation plate and transferred for single colony isolation of pGY177 and pGY178, respectively. As expected, no obvious phenotype was observed in isolated single colonies compared with the control. PCR genotyping indicated that four tested events from pGY177 exhibited PCR amplification, resulting in an editing efficiency of 20%, while only one event showed PCR amplification from pGY178, with an editing efficiency of 5% (Figure 6B and C). Sanger sequencing confirmed the large fragment deletions ranging in size from 102,228 to 102,249 bp in four edited events from pGY177 (Figure 6D). Interestingly, in addition to the large deletion, a 149 bp insertion was detected in event #1, and a 4 bp insertion was found in event #16. Additionally, a large sequence spanning 101,813 bp was excised via the Cas12a system in event #19 (Figure 6D).Taken together, these findings indicate that large genomic fragment deletions exceeding 100 kb can be efficiently achieved through CRISPR systems in *A*. *niger*.

**Figure 6.**
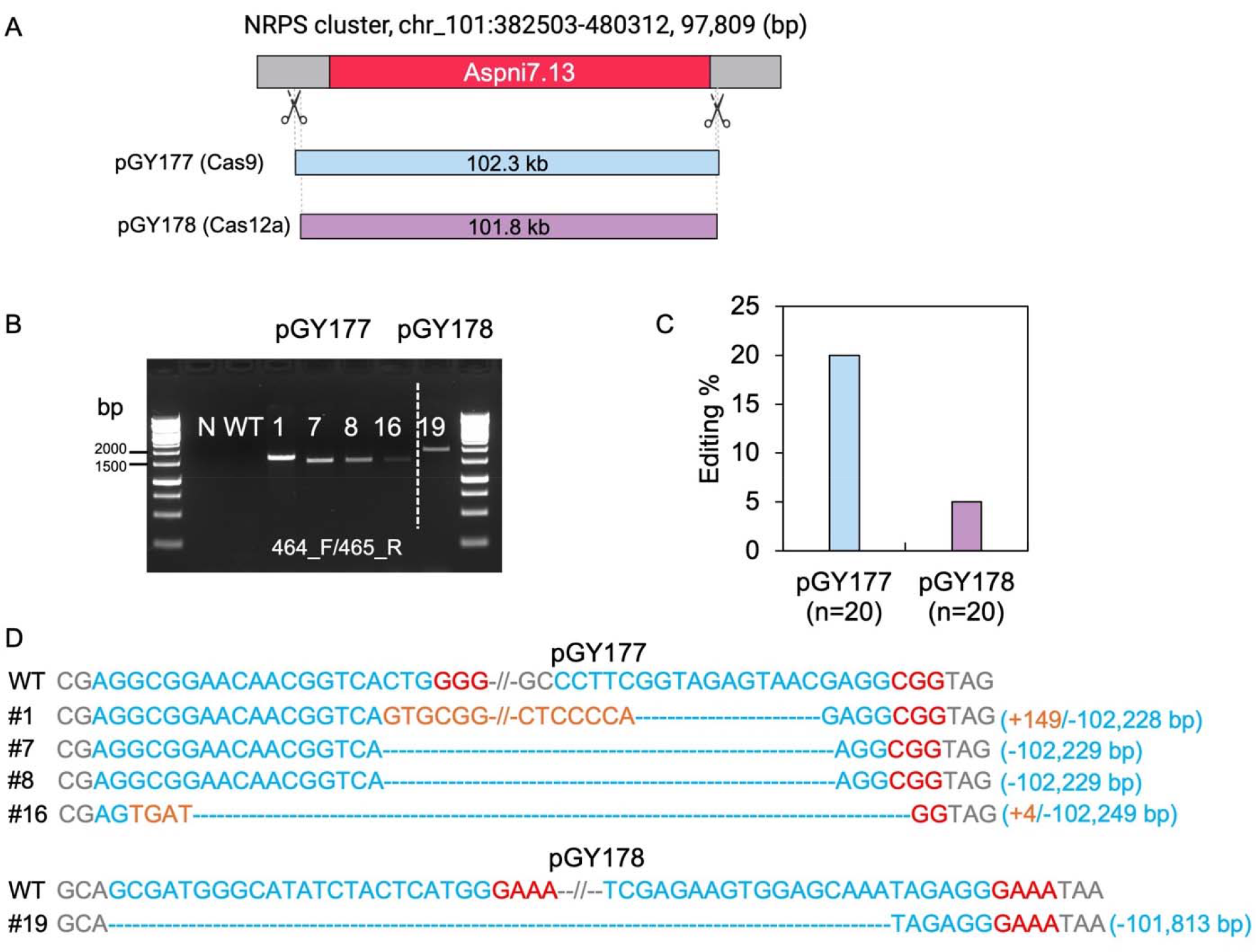
CRISPR-Cas9 and Cas12a mediated 100-kb deletion in *A. niger*. (A). Diagram of selected target region for 100-kb deletion. (B). PCR screening for mutants. N, water as negative control; WT, wild type as positive control. (C). Editing efficiency of 100-kb deletion. (D). Sanger sequencing for mutation confirmation. PAM sequences are highlighted in red, gRNA sequences in blue, and insertions are marked in orange.

## 4 Discussion

In this study, we have developed highly efficient, versatile, and programmable fungal CRISPR- Cas9 and Cas12a systems. Our research underscores their substantial potential for fungal genetic engineering, as evidenced by several key features: (i) rapid assembly of multiple gRNAs targeting different loci into a CRISPR-Cas9/Cas12a vector through a one-pot reaction; (ii) the critical role of endogenous tRNA^Ala^, tRNA^Phe^, tRNA^Arg^, tRNA^Ile^, and tRNA^Leu^ sequences in maintaining high gene-editing activity; (iii) both systems achieving up to 100% efficiency in single-gene editing; (iv) the superiority of CRISPR-Cas12a over Cas9, especially when employing a single RNA for gene disruption; (v) the Cas12a system’s advantage in attaining pure editing events, simplifying single spore isolation for pure fungal cultures; (vi) efficient induction of large-scale genomic deletions spanning at least 40 kb by both Cas9 and Cas12a systems; (vii) the capability of CRISPR systems to induce deletions at the 100-kb scale, albeit with reduced efficiency compared to smaller fragment deletions, yet still promising significant potential.

A significant advantage of CRISPR technology lies in its ease and efficiency in targeting multiple loci simultaneously, facilitated by multiplex CRISPR systems (McCarty et al. 2020). This approach involves incorporating multiple gRNAs into a single plasmid through molecular cloning. In this study, we developed two new vectors, pGY18 for the CRISPR-Cas9 system and pGY53 for the Cas12a system. These vectors include all critical components required for gene editing, except for the user-defined guide RNAs (gRNAs). pGY18 and pGY53 are designed to serve as vector backbones, enabling the construction of gene disruption vectors by simply inserting user-defined gRNAs via BsaI-based Golden Gate Assembly. While various cloning methods, such as SLIC, Gibson assembly, In-fusion, or SLiCE, enable the seamless joining of multiple fragments, they typically only allow the assembly of up to five or six fragments in one reaction (Li and Elledge 2007). Besides, two-step cloning is often required for the assembly of the final expression vector (McCarty et al. 2019; Hahn et al. 2020; Oh et al. 2020). However, recent advancements in comprehending ligase fidelity, bias, and efficiency in Golden Gate Assembly have enabled the assembly of over 50 fragments in a single reaction (Pryor et al. 2022). The BsaI-based Golden Gate Assembly has been demonstrated to facilitate the one-step assembly of multiple gRNAs into a CRISPR vector, accommodating up to 18 gRNAs (Yuan et al. 2022). Therefore, it is evident that the newly constructed vectors pGY18 and pGY53 can be readily programmed to target multiple loci in fungal genome editing.

Both Cas9 and Cas12a offer distinct advantages over conventional editing methods for fungal genome editing. While Cas9’s widespread use and well-understood mechanisms make it a popular choice, Cas12a’s capacity to recognize a T-rich PAM sequence significantly broadens the range of targetable genomic regions. Moreover, extensive comparisons between these two systems have been conducted across diverse species. For instance, studies have shown that Cas12a exhibits very low off-target effects in human cells and plants (Kim et al. 2016; Kleinstiver et al. 2016; Tang et al. 2018). Compared to the wildtype SpCas9 protein, the *Lachnospiraceae bacterium* Cas12a (LbCas12a) shows a higher editing efficiency in rice (Banakar et al. 2020). In tomatoes, comparable overall efficiencies were noted between SpCas9 and LbCas12a, with LbCas12a inducing more and larger deletions than SpCas9 (Slaman et al. 2024). In maize, Cas12a exhibited lower efficiency at the two target sites compared to the tested Cas9 system (Lee et al. 2019). In our study, we parallelly investigated the activities and specificities of CRISPR-Cas9 and Cas12a nucleases for targeted mutagenesis in filamentous fungi. Our findings revealed that Cas12a demonstrated superior efficiency compared to Cas9 when using a single gRNA for single-gene targeting. For Cas9-mediated single-gene editing, the use of double gRNAs is necessary to maintain high editing efficiency. Additionally, Cas12a tended to generate pure edited mutants, simplifying the process of isolating single spores required for screening and verifying fungal mutants. It’s worth noting that both our Cas9 and Cas12a systems utilize AMA1-based plasmids, facilitating plasmid loss through subculturing in a nonselective cultures (Aleksenko and Clutterbuck 1997). This enables marker-free gene editing and recyclable usage of the same marker, while also reducing the risk of potential off-target effects in the resulted strains.

To facilitate strain engineering and laboratory evolution of various host strains, numerous tools for large fragment deletion have been developed. For example, while double-stranded DNA breaks directed by CRISPR are often highly lethal in many bacteria, Standage-Beier et al. demonstrated that dual-targeted nicking allows for the deletion of 133 kb of the genome in *Escherichia coli* (Standage-Beier et al. 2015). Li et al. described the rapid deletion of a large DNA fragment of approximately 38 kb between the two genes *TRM10* and *REX4* using CRISPR/Cpf1 in *Saccharomyces cerevisiae* (Li et al. 2018). Large DNA fragment deletion in filamentous fungi has also been explored across various species, with success often dependent on the utilization of selection markers and homology-directed repair templates (donor DNAs). For instance, large chromosomal deletions in koji molds, *A*. *oryzae* and *A*. *sojae*, reaching up to 100 kb and 200 kb respectively, rely on NHEJ-deficient Δ*ku70* strains for their success, while achieving multi-segment deletions entails a cumbersome procedure involving 5-fluoroorotic acid to remove the selectable marker (pyrG) and restore auxotrophy (Takahashi et al. 2008). Using homology-directed repair templates, large DNA fragments up to 31.5 kb involved in the biosynthesis of the yellow compound sorbicillinoid were effectively deleted via the CRISPR- Cas9 system in *Acremonium chrysogenum* (Chen et al. 2020). A 10 kb deletion of a metabolic synthetic cluster was accomplished using the CRISPR-Cas9-TRAMA system in conjunction with a homologous template in *Cordyceps militaris* (Chen et al. 2022). Most recently, chromosomal segments of 201 kb and 301 kb were effectively deleted in *Aspergillus flavus* using a dual CRISPR/Cas9 system, achieving deletion efficiencies ranging from 57.7% to 69.2%, respectively (Chang 2024). The deletion efficiencies showed significant improvement in the presence of a single copy of donor DNA compared to samples lacking donor DNA. However, the utilization of CRISPR-Cas12a for large fragment deletion in fungi is rarely explored. In this study, we demonstrate that both CRISPR-Cas9 and Cas12a are capable of efficiently inducing large fragment deletions of varying sizes in *A*. *niger*, reaching up to 102 kb, which accounts for 0.3% of the *A*. *niger* genome, with remarkable efficiency of 20%. Unlike those reported systems that necessitate the expression of multiple constructs and the addition of donor DNA, our systems only require the introduction of a single plasmid, rendering them the simplest and most straightforward to date. Moreover, the re-ligation of genomic endpoints resulting from DSBs is primarily repaired by NHEJ. Further investigation is necessary to explore the potential of these systems in inducing larger fragment deletions exceeding 100 kb and enhancing their deletion efficiency. Additionally, evaluating the impact of incorporating homology-directed repair templates in these systems could be advantageous for inducing larger deletions.

In conclusion, we have developed a suite of high-efficiency CRISPR systems for marker-free gene disruption to expand the CRISPR toolkit and accelerate metabolic engineering in fungi for enhancing the production of biofuels and bioproducts. These multiplex CRISPR systems enable researchers to quickly target multiple genetic loci simultaneously by incorporating multiple gRNAs into a single plasmid through cloning. The remarkable efficiency of these CRISPR systems will streamline genetic engineering processes, facilitating their application not only in model fungal species but also in non-model species. Additionally, by eliminating the requirement of selectable markers, these systems reduce the complexity and increase the speed of genetic manipulations, making them more accessible and practical for a wide range of fungal studies.

## Data availability statement

The data generated or analyzed during this study are included within this published article and its supplementary information files. The constructs will be available via Addgene (https://www.addgene.org/).

## Author contributions

GY: data curation, formal analysis, investigation, methodology, validation, visualization, writing–original draft, writing–review and editing. SD: supervision, resources, investigation and writing–review and editing. JC: writing–review and editing and investigation. ZD: writing– review and editing and investigation. BH: project administration, resources and writing–review and editing. JK: writing–review and editing and investigation. KP: writing–review and editing, supervision, investigation, funding acquisition, and conceptualization.

---

**Supplementary Figure S1.**
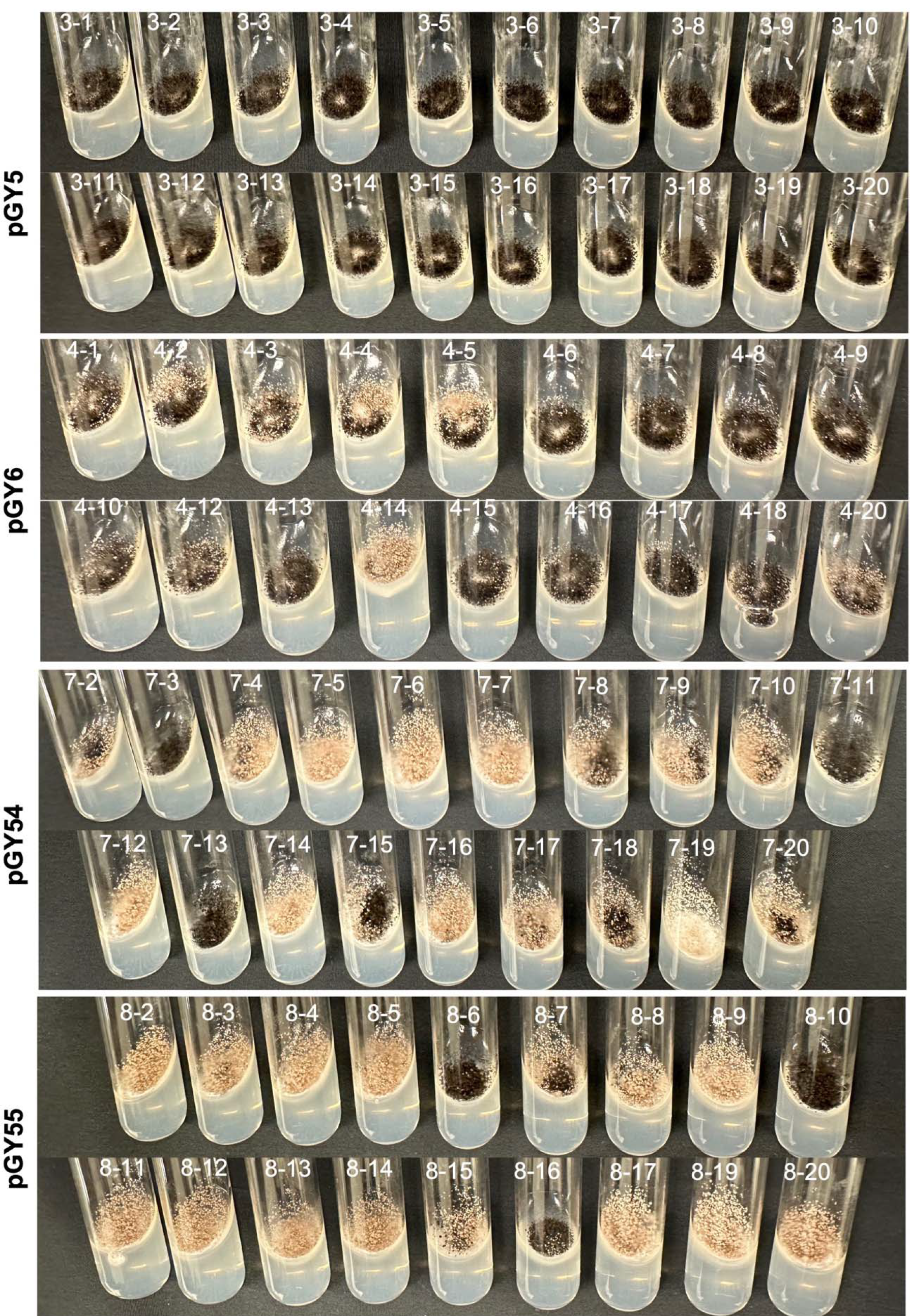
**Phenotypic characteristics of transformants targeting the *albA* gene after single colony isolation on an minimal medium agar slant.**

**Supplementary Figure S2.**
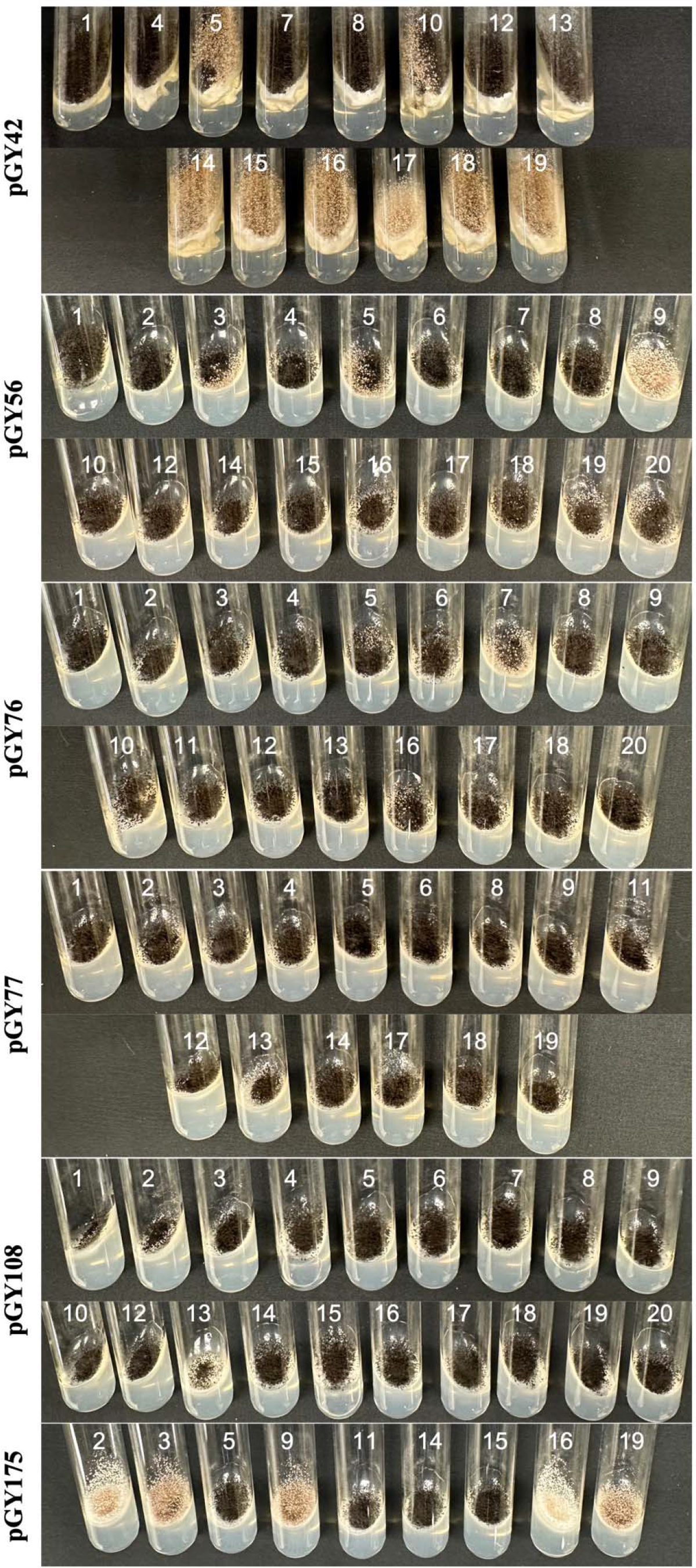
**Phenotypic characteristics of selected transformants with larger fragment deletions after isolating single colonies on an agar slant.**

